## Supplementary material for "The Drosophila EGF domain protein Uninflatable sets the switch between wrapping glia growth and axon wrapping instructed by Notch": Materials and Methods

**Key resources table**

| REAGENT or RESOURCE | SOURCE | IDENTIFIER |
| --- | --- | --- |
| Antibodies | | |
| anti-dsRed 1:1000 | Clontech Labs 3P | 632496 |
| anti-LacZ 1:10 | Hybridoma | 40-1a |
| anti-Repo 1:5 | Developmental Studies Hybridoma Bank | 8D12 |
| anti-HRP-DyLight^TM^649 1:500 | Dianova | 123-165-021 |
| anti-GFP 1:1000 | Invitrogen | A6455 |
| anti-mouse 488 | Invitrogen | A10680 |
| anti-mouse 568 | Invitrogen | A11031 |
| anti-rabbit 488 | Invitrogen | A11008 |
| anti-rabbit 568 | Invitrogen | A11011 |
| Bacterial and virus strains | | |
| E. coli OneShot^®^ TOP10 chemically competent cells | Invitrogen | C404003 |
| Experimental models: Organisms/strains | | |
| w^1118^ | Bloomington Drosophila Stock Center | BDSC:3605 |
| N^[ts]1^ | [Shellenbarger, 1975 #5642] |  |
| UAS-LacZ^NLS^ | [Hummel, 2002 #5641] |  |
| UAS-H2B-mRFP | Bloomington Drosophila Stock Center | BDSC:94270 |
| UAS-CD8::mCherry | Bloomington Drosophila Stock Center | BDSC:27391 |
| UAS-λhtl | A. Michelson |  |
| UAS-htl^DN^ | Bloomington Drosophila Stock Center | BDSC:5366 |
| UAS-N^ICD^ | T. Klein |  |
| UAS-uif::GFP | M. Gonzalez-Gaitan | [Loubéry, 2014 #2770] |
| UAS-GFP^dsRNA^ | Bloomington Drosophila Stock Center | BDSC:9331 |
| UAS-mCherry^dsRNA^ | Bloomington Drosophila Stock Center | BDSC:35785 |
| UAS-LacZ^dsRNA^ | S. Schirmeier |  |
| UAS-uif^dsRNA HMS01822^ | Bloomington Drosophila Stock Center | BDSC:38354 |
| UAS-N^dsRNA GD144^ | Vienna Drosophila Resource Center | VDRC1112 |
| UAS-N^dsRNA 9G^ | Bloomington Drosophila Stock Center | BDSC:7077 |
| UAS-Dl^dsRNA GD2642^ | VDRC | VDRC37287 |
| U6-Dl^sgRNA^ |  | BDSC:83095 |
| UAS-Ser^dsRNA^ ^GD14442^ | VDRC | VDRC27172 |
| U6-Ser^sgRNS^ |  | BDSC:84169 |
| UAS-Su(H)^dsRNA HMS05748^ | Bloomington Drosophila Stock Center | BDSC:67928 |
| UAS-mam^dsRNA KK110687^ | Vienna Drosophila Resource Center | VDRC102091 |
| UAS-Cont^dsRNA GD12610^ | Vienna Drosophila Resource Center | VDRC28294 |
| UAS-Cont^dsRNA GD12610^ | Vienna Drosophila Resource Center | VDRC40613 |
| UAS-Cas9 | Bloomington Drosophila Stock Center | BDSC:58985 |
| GFP^sgRNA^ | S. Schirmeier |  |
| uif^sgRNA 2nd Exon^ | This work |  |
| uif^sgRNA CS^ | This work |  |
| uif^sgRNA TMD^ | This work |  |
| uif^sgRNA CD^ | This work |  |
| N^sgRNA^ | Bloomington Drosophila Stock Center | BDSC:84168 |
| Dl^MI04868-TG4.1^ | Bloomington Drosophila Stock Center | BDSC:77753 |
| nrv2::GFP | Bloomington Drosophila Stock Center | BDSC:6828 |
| nSyb-Gal4 | Bloomington Drosophila Stock Center | BDSC:51635 |
| nrv2-Gal4;R90C03-Gal80,UAS-CD8::Cherry | [Kottmeier, 2020 #2674] |  |
| Gbe+Su(H)-lacZ | [Furriols, 2001 #5328] |  |
| repo4.3-stg-GFP | This paper |  |
| nrv2-Gal4 | Bloomington Drosophila Stock Center | BDSC:6800 |
| Oligonucleotides | | |
| sgRNA for *uif* cleavage just before transmembrane domain: Uif2ndExon_gRNA2_fw: GTCGTTTCAATATCAAGCACTCGT | This paper | N/A |
| sgRNA for *uif* cleavage just before transmembrane domain: Uif2ndExon_gRNA2_rev: AAACACGAGTGCTTGATATTGAAA | This paper | N/A |
| sgRNA for *uif* cleavage just before transmembrane domain: UifCS_gRNA2_fw: GTCGTGTTCTGCGTACCTCGGTAG | This paper | N/A |
| sgRNA for *uif* cleavage just before transmembrane domain: UifCS_gRNA2_rev: AAATCTACCGAGGTACGCAGAACA | This paper | N/A |
| sgRNA for *uif* cleavage just before transmembrane domain: UifTMD_gRNA4_fw: GTCGCGCTGTGTGGGCTCCTTTAC | This paper | N/A |
| sgRNA for *uif* cleavage just before transmembrane domain: UifTMD_gRNA4_rev: AAACGTAAAGGAGCCCACACAGCG | This paper | N/A |
| sgRNA for *uif* cleavage just before transmembrane domain: UifCD_gRNA1_fw: GTCGCTACAATGAAACGTACATGA | This paper | N/A |
| sgRNA for *uif* cleavage just before transmembrane domain: UifCD_gRNA1_rev: AAACTCATGTACGTTTCATTGTAG | This paper | N/A |
| Recombinant DNA | | |
| pUAST-dU63gRNA vector carrying a ubiquitous U6:3 promoter | S. Schirmeier |  |
| Software and algorithms | | |
| Fiji | [Schindelin, 2012 #1257] |  |
