## Supplementary Figures for "The Drosophila EGF domain protein Uninflatable sets the switch between wrapping glia growth and axon wrapping instructed by Notch"

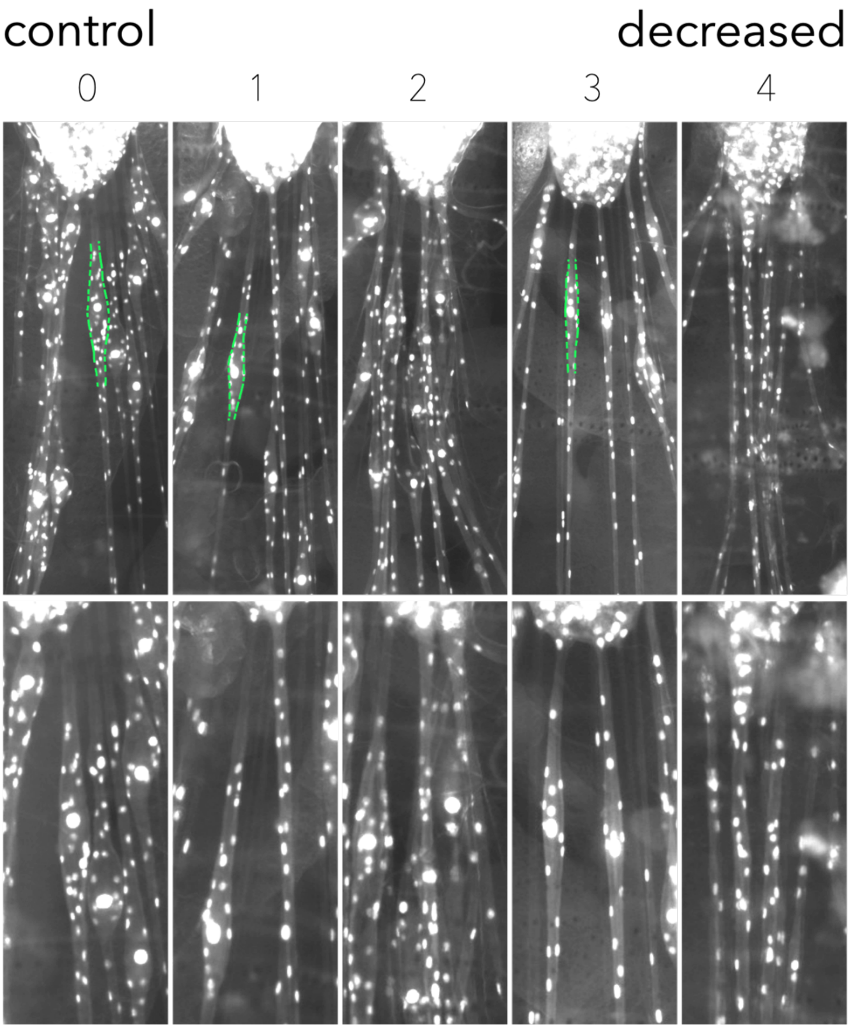


**Figure S1 Suppressor screen for genes downstream of activated *heartless,* related to Figure 1.** Exemplary images of living third instar larvae expressing *repo-stinger::GFP*. Analyzed genes were classified according the changes in bulge size. 0 = no change; 1, 2 = slight changes in the bulge size. 3 = strong suppression of the bulge size; 4 = complete suppression of the bulging phenotype. For list of genes screened, please see supplementary Table 1.

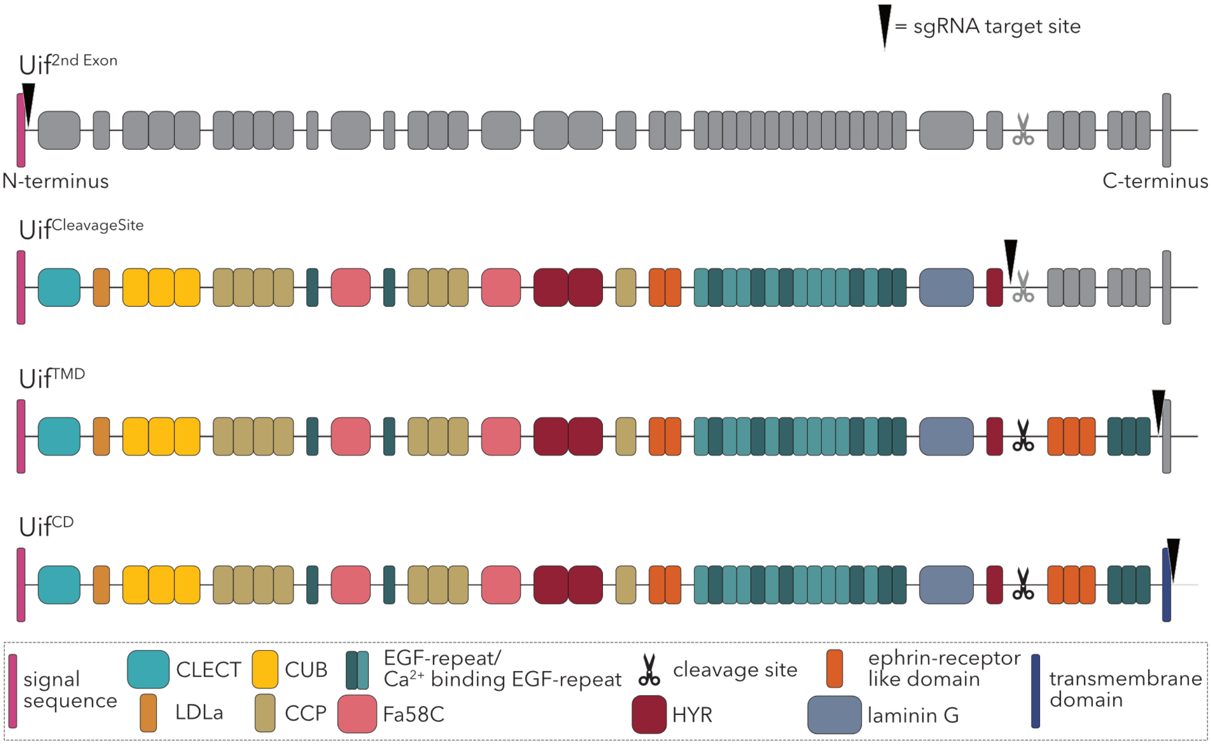


**Figure S2 Schematic representation of sgRNA target sites used for CRISPR mediated *uif* knockout, related to Figure 2.** Target sites (black arrowhead) and predicted protein residues (Uif isoform B; present domains shown in color, absent domains shown in grey). Sequence of domains from N- to C-terminus: signal sequence (pink); one C-type lectin-like (CLECT) domain (turquoise), proposed to be a calcium-dependent carbohydrate binding module (Cambi & Figdor, 2003); one low-density lipoprotein receptor protein class A (LDLa) domain (dark yellow), predicted to have a protein binding function (May et al., 2007); three CUB domains (light yellow), a structural domain of unknown function often involved in developmental processes (Bork & Beckmann, 1993); eight CCP domains (beige), predicted to be adhesion protein interaction domains (Norman et al., 1990; Reid & Day, 1989); 18 EGF-like-repeats (10 are $Ca^{2+}$ binding; light/dark petrol), which are protein interaction domains involved in cell signaling (Appella et al., 1988); two coagulation factor 5/8 C-terminal (FA58C) domains (light red), which are carbohydrate binding motifs (L. Zhang & Ward, 2009); three hyaline repeat (HYR) domains (dark red), with a putative function in cell adhesion (Carroll et al., 2008); five ephrin-like domains (orange) and one laminin G domain (light blue), also likely involved in cell adhesion and signaling (Banerjee et al., 2011; Deutzmann et al., 1988); proteolytic cleavage site (scissors) at amino acid 2991 (L. Zhang & Ward, 2009); antibody binding site (AB) at amino acid 2882-3157 (L. Zhang & Ward, 2009); additional exon isoform C indicated by red line; transmembrane domain (dark blue); intracellular C-terminus does not contain any conserved domains (L. Zhang & Ward, 2009).

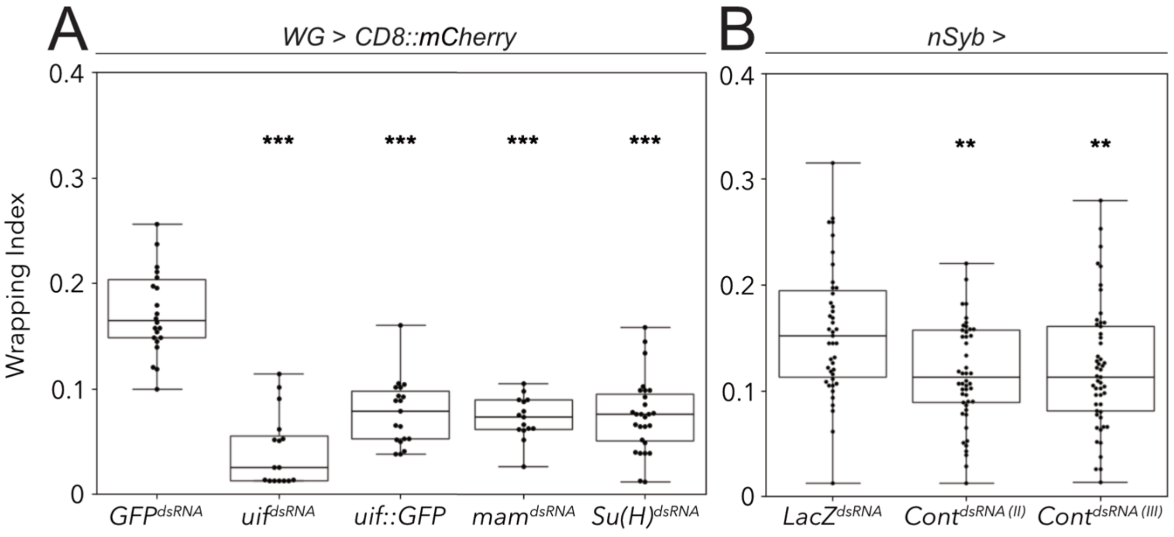


**Figure S3 Quantification of the wrapping index, related to Figure 3, 5, 6.** Wrapping Index (WI) quantified from electron microscopic cross sections of third instar larval abdominal peripheral nerves with (**A**) wrapping glia specific (*nrv2-Gal4;R90C03-Gal80*) expression of mock control *GFP*$^{dsRNA}$ (WI=0.17), *uif* dsRNA$^{HMS01822}$, *uif::GFP, Su(H)*$^{dsRNA}$ and *mam*$^{dsRNA}$ is plotted. Both *uif* knockdown (WI=0.03) and overexpression (WI=0.08) lead to a significant reduction of axonal wrapping (comparing the WI of control and *uif*$^{dsRNA}$ larvae: $p=2.88\text{x}10^{-7}$; comparing the WI of control and *uif::GFP* larvae: $p=2.56\text{x}10^{-9}$). Similarly, knockdown of the Notch downstream components *Su(H)* and *mam* reduces the wrapping index significantly (*mam*$^{dsRNA}$, WI=0.07, $p=4.29\text{x}10^{-9}$; *Su(H)*$^{dsRNA}$, WI=0.075, $p=4.21\text{x}10^{-11}$). (B) Neuronal knockdown (*nSyb-Gal4*) of *Contactin* leads to a significant reduction of the wrapping index from 0.15 to 0.11 (for both RNAi lines p=0,006). For statistical analysis a t-test was performed for normally distributed data (Shapiro-test), a Mann-Whitney-U test was performed for not normally distributed data. *GFP*$^{dsRNA}$ n=4 larvae with 4-7 nerves per specimen; *uif*$^{dsRNA}$ n=3 larvae, 5-9 nerves per specimen; *uif::GFP* n=3 larvae, 5-6 nerves per specimen; *mam*$^{dsRNA}$ n=3 larvae with 5-9 nerves per specimen; *Su(H)*$^{dsRNA}$ n=4 larvae with 6-8 nerves per specimen; *Cont*$^{dsRNA}$ (both) n=5 larvae with 10 nerves per specimen. □=0.05, ** p≤0.01, *** p≤0.001.

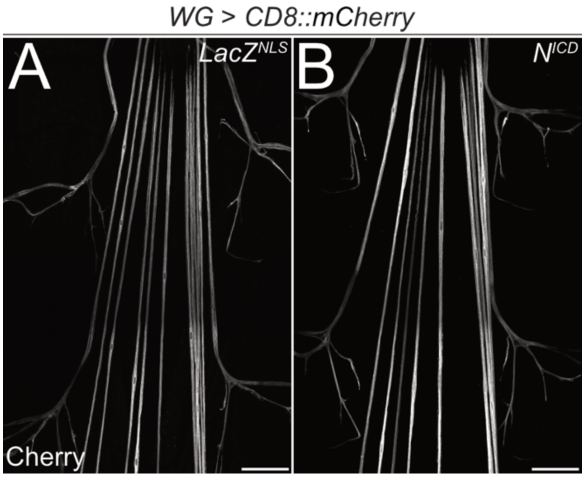


**Figure S4 Notch activation impairs wrapping glial development at higher temperatures, related to Figure 4.** Filet preparations of wandering third instar larvae stained for CD8::mCherry expression. (**A**) Nuclear *LacZ* as mock control for (**B**) overexpression of Notch intracellular domain in wrapping glia specifically (*nrv2-Gal4;R90C03-Gal80*). $N^{ICD}$ overexpression does not affect wrapping glial morphology. n=5 larvae for all genotypes. Scale bars 100 μm.

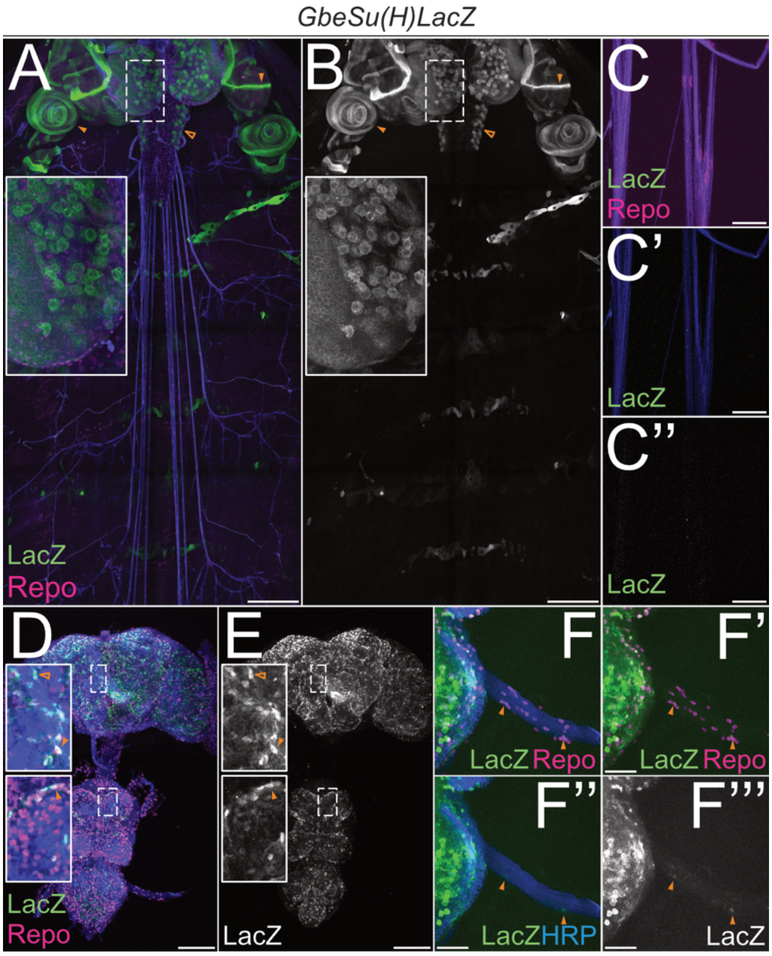


**Figure S5 The Notch activity reporter is active in differentiated glia, related to Figure 4, 5.** Maximum intensity projection of a confocal image stack of third instar larval filet and adult brain preparations stained for anti-LacZ, representing Notch activity (green), glial nuclei (anti-Repo, magenta) and neuronal membranes (anti-HRP, blue). Bacterial *LacZ* gene under the control of three copies of palindromic *Grainyhead* (*Grh*) binding sites (*Gbe*) and *Su(H)* binding is active specifically in cells where Notch signaling is present. (**A**, **B**) Expression can be detected in neuroblasts of the brain lobes (asterisk and close up) and the thoracic neuromeres (unfilled arrowhead). Additional Notch activity is detected in the imaginal discs (filled arrowheads); scale bar 200 µm. (**C**-**C''**) No Notch activity is found in larval peripheral glial nuclei; scale bar 100 µm. (**D**, **E**) LacZ staining overlaps with Repo staining, indicating expression in adult glia (close ups, filled arrowheads) which is also detectable in non-glial cells (close ups, unfilled arrowheads). Not all Repo positive cells show LacZ staining (close ups, asterisk), indicating that Notch signaling is not active in all glia. Scale bar 100 µm. (**F**-**F'''**) Single plane of Z-stack is shown. Some peripheral glial cells show LacZ staining (arrowheads); scale bar 100 µm. n=3 animals for both developmental stages.

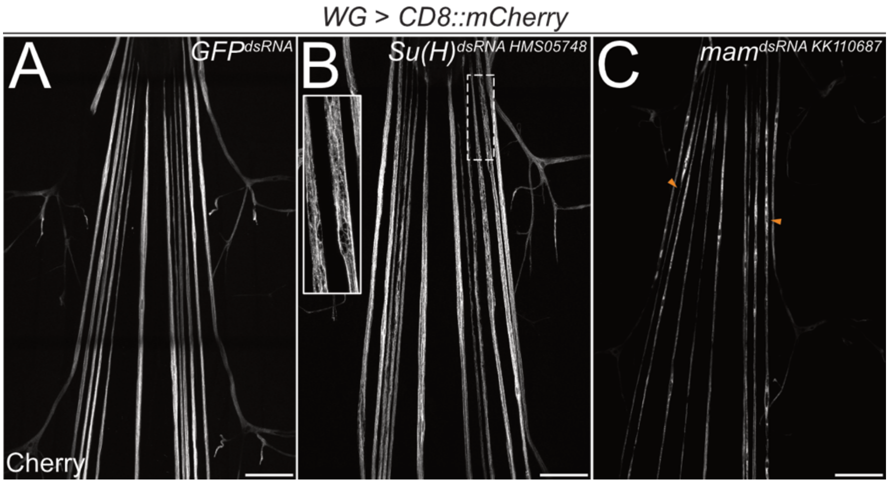


**Figure S6 Knockdown of *mam* and *Su(H)* impairs wrapping glial development, related to Figure 5.** Filet preparations of wandering third instar larvae stained for CD8::mCherry expression. (**A**) Wrapping glia specific (*nrv2-Gal4;R90C03-Gal80*) expression of *GFP*$^{dsRNA}$ as mock control, (**B**) *mastermind* (*mam*) *dsRNA*$^{KK110687}$, or (**C**) *Suppressor of Hairless* (*Su(H)*) *dsRNA*$^{HMS05748}$. Note impaired wrapping glial morphology (B, C, arrowheads). n=5 larvae for all genotypes. Scale bars 100 µm.

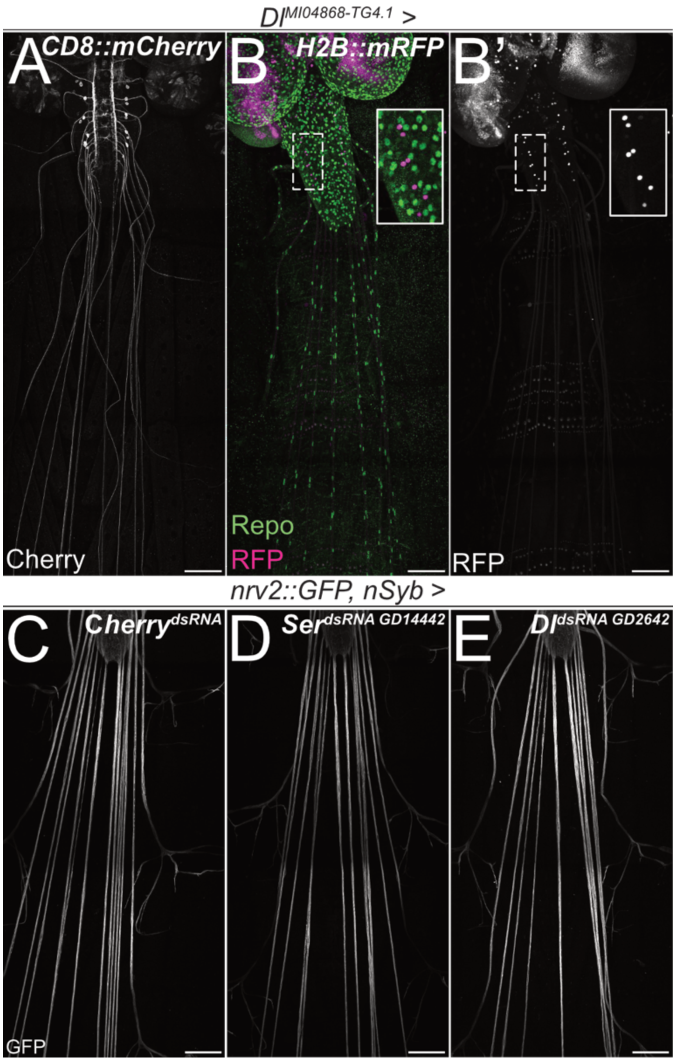


**Figure S7 Neuronal knockdown of Notch ligands *Dl* and *Ser* does not affect wrapping glial morphology, related to Figures 3, 4.** Filet preparations of wandering third instar larvae stained for (**A**) CD8::mCherry expression or (**B**) Repo (stains glial nuclei, green) and H2B-mRFP expression. A Trojan-Gal4 insertion in the *Delta* (*Dl*) locus [$Dl^{MI04868\text{-}TG4.1}$] drives expression of (A) *CD8::mCherry* targeting membranes or (B) *H2B::mRFP* targeting nuclei. Two *Dl* expressing neurons are found in each hemineuromere and show no *repo* expression (B and B', close up). n=5 larvae for all genotypes. Scale bars 100 µm. (**C-E**) Filet preparations stained for *nrv2::GFP* expression. (C) Neuron specific expression of mock control C*herry* dsRNA (D) *Serrate* (*Ser*) $dsRNA^{GD14442}$ and (E) *Delta* (*Dl*) $dsRNA^{GD2642}$. Neither *Ser* nor *Dl* knockdown in neurons affects wrapping glial morphology. n=5 larvae for all genotypes. Scale bars 100 µm.

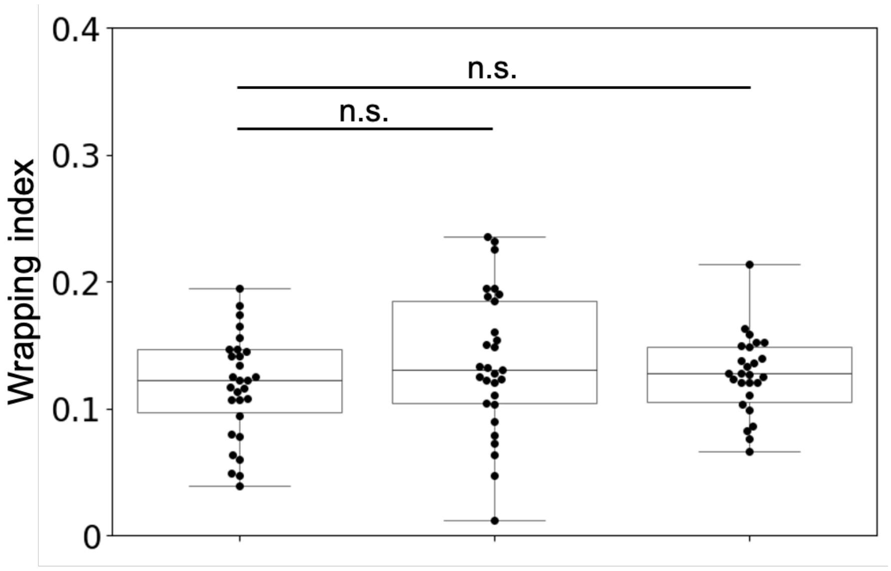


| | nSyb >tdTomato,Cas9 | nSyb > tdTomato, Cas9, Delta-sgRNA | nSyb > tdTomato, Cas9, Serrate-sgRNA |
|---|---|---|---|
| Animals | 3 | 3 | 3 |
| Nerves | 30 | 29 | 30 |
| Median wrapping index | 0.122 | 0.13 | 0.1275 |
| Shapiro | 0.82 | 0.8 | 0.0003 |
| p-value | 0.24 (t-test) | | - |
| | 0.56 (MW-U-test) | | |

**Figure S8 Neuronal knockout of Notch ligands *Dl* and *Ser* does not affect wrapping glial morphology, related to Figures 3, 4** Filet preparations of wandering third instar larvae with the genotypes indicated were sectioned 150µm distant from the tip of the ventral nerve cord. Electron microscopic images were analyzed for their wrapping index. Cas9 mediated neuronal knockout of neither Delta nor Serrate significantly affects the wrapping index.

**Table S1- S3,**

**Figure source data**

Please see attached Excel files
